# Transcriptional read-through of the long non-coding RNA *SVALKA* governs plant cold acclimation

**DOI:** 10.1101/287946

**Authors:** Peter Kindgren, Ryan Ard, Maxim Ivanov, Sebastian Marquardt

**Affiliations:** University of Copenhagen, Department of Plant and Environmental Sciences, Copenhagen Plant Science Centre, Frederiksberg, Denmark

## Abstract

Most DNA in the genomes of higher organisms does not encode proteins, but is transcribed by RNA polymerase II (RNAPII) into long non-coding RNA (lncRNA). The biological significance of most lncRNA is largely unclear. Here, we identify a lncRNA (*SVALKA*) in a cold-sensitive region of the *Arabidopsis* genome. Mutations in *SVALKA* affect the timing of maximal *CBF1* expression and freezing tolerance. RNAPII read-through transcription of *SVALKA* results in a cryptic lncRNA overlapping *CBF1* on the antisense strand, termed *asCBF1*. *asCBF1* transcription is anti-correlated with *CBF1* expression. Our molecular dissection reveals that *CBF1* is suppressed by RNAPII collision stemming from the *SVALKA-asCBF1* lncRNA cascade. The *SVK-asCBF1* cascade provides a mechanism to tightly control *CBF1* expression and timing that could be exploited to maximize freezing tolerance with mitigated fitness costs. Inversion of the transcriptional direction of a lncRNA cascade relative to the genes in a co-regulated cluster provides an elegant inbuilt negative feedback for cluster expression. Our results provide a compelling example of local gene regulation by lncRNA transcription having a profound impact on the ability of plants to appropriately acclimate to suboptimal environmental conditions.

## Main

RNA Polymerase II (RNAPII) transcription in genomes results in the production of many long non-coding RNAs (lncRNAs)^1^. The functional significance of resulting lncRNA molecules is actively debated even though biological roles are identified for an increasing number of examples^2–4^. Expression of lncRNAs is remarkably specific to the environmental condition, tissue or cell type, arguing for roles of lncRNA in regulation^5–7^. In addition to functions carried out by lncRNA molecules, the process of transcribing non-coding DNA sequences can by itself be regulatory in many systems^8,9^. Non-coding DNA regions in genomes can therefore affect gene expression by different mechanisms that need to be resolved experimentally.

Sessile organisms respond to changing environmental conditions with regulation of gene expression. Early events in the *Arabidopsis* cold response include rapid transcriptional up-regulation of the intron-less C-repeat/dehydration-responsive element binding factors (CBFs)^10,11^. The CBFs are highly conserved transcription factors that promote cold tolerance in many plant species and are often arranged in a single cluster^12^. *CBF* expression during cold exposure activates downstream *COLD REGULATED (COR)* genes that promote freezing tolerance by adjusting the physiological and biochemical properties of plant cell interiors^13–15^. Intriguingly, constitutive *CBF* expression increases cold-tolerance but is also associated with fitness penalties^16–19^. The expression of the endogenous *CBF1* gene is characterized by a transient peak of maximal expression^11^. Potential roles of non-coding transcription during *CBF* induction and repression are currently unclear.

The plant response following long exposure to cold temperatures is associated with numerous lncRNA that associate with the *FLOWERING LOCUS C* (*FLC*) gene. The antisense lncRNA *COOLAIR* is induced after 2 weeks of cold, and initiates chromatin repression of the *FLC* gene^2,20^. Full *FLC* silencing is aided by polycomb repressive complex recruitment by the intron-derived lncRNA *COLDAIR* and reinforced by the promoter derived lncRNA *COLDWRAP^21^*. This cold-triggered cascade of lncRNA establishes long-lasting and stable repression of the *FLC* gene and serves as a paradigm for gene repression by lncRNA^22^. However, it is currently unresolved if early responses to cold temperatures in plants are mediated by lncRNA.

Here, we map transcriptional start sites in *Arabidopsis* during early responses to cold temperatures. We identify a cascade of two lncRNA that fine-tune the expression of the *CBF1* gene. Transcriptional read-through of the lncRNA *SVALKA* results in the expression of a cryptic antisense *CBF1* lncRNA (*asCBF1*). *asCBF1* transcription results in RNAPII collision to limit the expression of full length *CBF1*. This work extends the biological roles of lncRNA during cold acclimation and it provides an elegant mechanism that achieves rapid dynamic regulation of environmentally sensitive gene expression.

## Results

### Identification of the lncRNA SVALKA

To identify transcription initiation events that respond to cold temperature in *Arabidopsis*, we performed Transcription Start Site (TSS)-sequencing. Our analyses revealed 489 down-regulated and 1404 up-regulated TSSs in response to 3 hours at 4°C (Fig. 1a, Supplementary table 1). Most differentially used TSSs were located in promoter regions corresponding to changed promoter usage in fluctuating environments. Of the up-regulated genes (±200 bp of major TSS), the *CBF* genes were upregulated 100-400 fold making the *CBF* genomic region by far the most cold-responsive region in the genome (Fig. 1b). Interestingly, we detected a cold-responsive long non-coding RNA (lncRNA), transcribed on the antisense strand between *CBF3* and *CBF1* (Fig. 1c) that we named *SVALKA* (*SVK*). We found two TSSs by Rapid Amplification of cDNA Ends (RACE) that corresponded to the peaks we observed with TSS-seq (Fig. 1d). Our 3’RACE analysis also supported the existence of alternative *SVK* isoforms. Notably, we detected no transcript overlapping the *CBF1-3’UTR*. RT-qPCR revealed that more *SVK* is generated from the proximal promoter than from the distal promoter (Supplementary Fig. 1), consistent with TSS-seq analyses. A time series of cold exposure (4°C) with samples taken every 4 hours revealed the expression pattern of *SVK* and *CBF1* (Fig. 1e). *CBF1* expression peaked at 4 hours followed by a decrease that reached a relatively stable level after 12-16 hours, a pattern in congruence with earlier studies^23^. In contrast, *SVK* showed gradual increase in expression from 4 hours to reach a stable maximum level after 12-16 hours.

**Figure 1.**
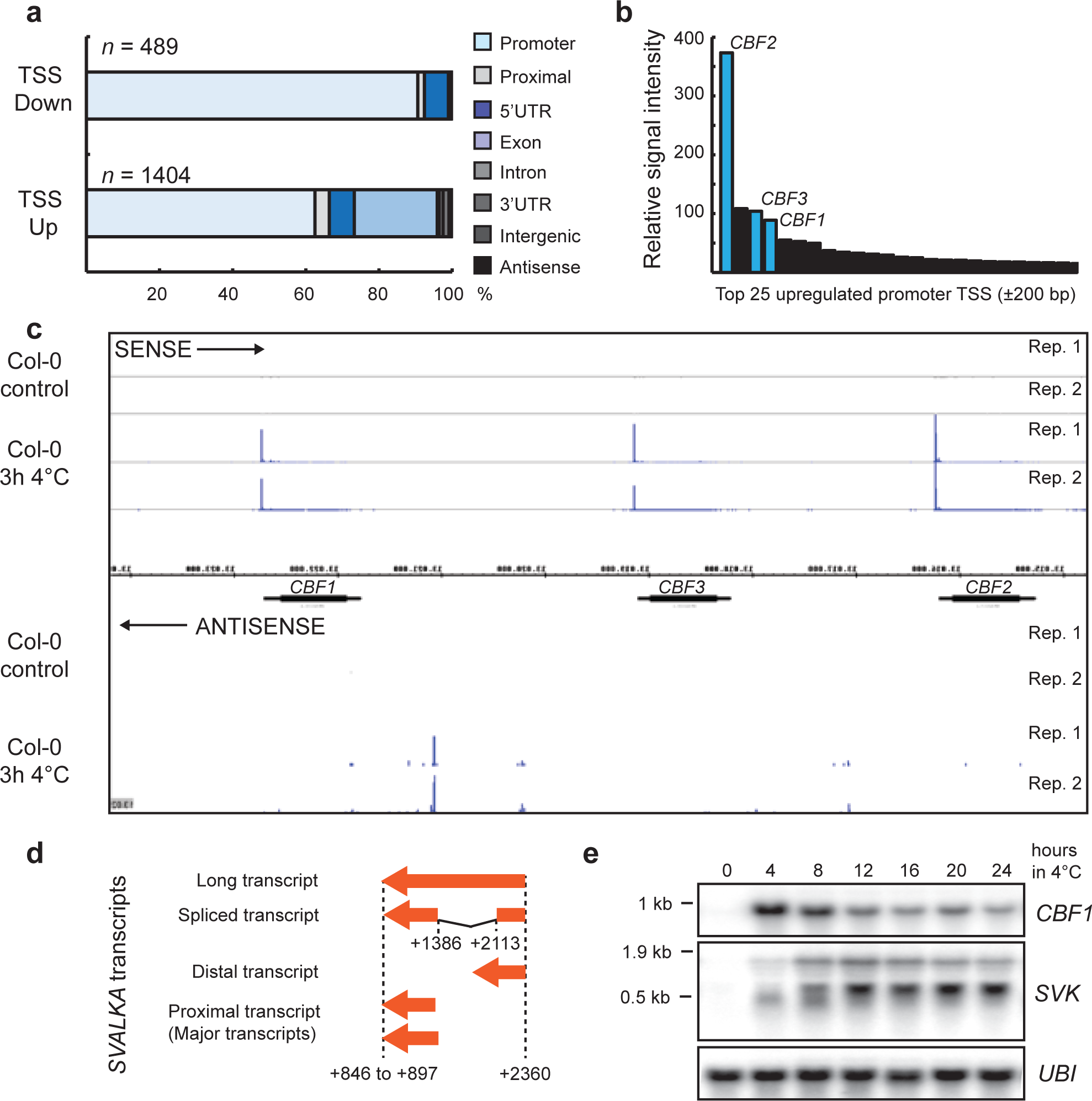
Transcription Start Site (TSS) Sequencing in Col-0 after 3 hours of exposure to 4°C identified the long non-coding RNA, *SVALKA*. a) Differentially expressed TSS after cold exposure in Col-0. A total of 1893 TSS changed their expression significantly (p<0.05). Of these, 489 were down-regulated while 1404 were up-regulated. The TSS were classified according to their position in the genome. The majority of differentially expressed TSS were found in or around promoters. b) Activity of up-regulated promoters after cold exposure identities the *CBF* genes (indicated in red bars) as highly up-regulated. Graph represents the top 25 of the up-regulated promoters in Col-0 after cold exposure. c) Screenshot of the *CBF* genomic region and the identified TSS. The upper panel shows the sense direction and clear TSS can be found for the three *CBF* genes. The lower panel shows the antisense direction where a group of cold induced TSS was identified in the intergenic region between *CBF1* and *CBF3*. d) Summary of 3’- and 5’RACE of *SVALKA*. Two clusters of capped TSS were found for *SVALKA*, a distal TSS centered around 2360 bp and a proximal TSS centered around 1386 bp in respect to the translation start of *CBF1*. A cluster of polyadenylation sites were found 846 to 897 bp in respect to the translation start of *CBF1*. The plethora of *SVALKA* transcripts includes a spliced isoform with a splice site from 1386 to 2113 bp in respect to the translation start of *CBF1*. e) RNA levels of *CBF1* and *SVALKA* during a time course of cold exposure of Col-0. While *CBF1* transiently is upregulated early in the cold response, *SVALKA* responds later and its expression is anti-correlated to sense *CBF1*. *UBI* is used as a loading control. Experiments were done with three biological replicates showing similar results.

### *SVK* represses *CBF1* and promotes cold acclimation

The anti-correlation between *CBF1* and *SVK* and their genomic proximity suggested that *SVK* could be involved in *CBF1* repression. To test this, we constructed two distinct *LUCIFERASE (LUC)* reporter lines of *CBF1* with different termination sequences. We expected any repressive role of *SVK* to be present in the lines with endogenous 3’-sequences (*CBF1*:*SVK)*. In contrast, we expected effects of *SVK* to be absent in the lines with the terminator sequences of the *NOPALINE SYNTHASE (NOS)* gene devoid of antisense transcription^4^ (*CBF1*:T_NOS_). We subjected three independent lines from each construct to 4°C for 0-12 hours and subsequently measured LUC activity (Fig. 2a-b). The three *CBF1*:T_NOS_ lines showed increased *LUC* activity compared to the *CBF1*:*SVK* lines. These data demonstrated that *SVK* represses *LUC* activity. We isolated two T-DNA lines that disrupted *SVK* (*svk-1*) or uncoupled *SVK* from *CBF1* (*uncoupling svalka-1, uns-1*) (Fig. 2c). *svk-1* showed non-detectable expression of *SVK* while the *uns-1* mutants showed wild-type levels of *SVK*. Interestingly, we detected *CBF1* mis-regulation in both mutants (Fig. 2d-e). *SVK* expression in *uns-1* argues against a trans-acting function for this lncRNA (Fig. 2d). The mutants showed an increase of *CBF1* response to cold exposure, supporting the notion that *SVK* plays a repressive role on *CBF1* mRNA levels. Furthermore, the effect was transmitted to the expression of CBF-activated *COR* genes (Fig. 2f). These findings suggested that *SVK* transcription could affect cold acclimation and freezing tolerance. We conducted a freeze test of wild-type, *svk-1* and *uns-1* plants to test the role of *SVK* in freezing tolerance (Fig. 2g). Cold-acclimated *svk-1* and *uns-1* showed a higher freezing tolerance compared to wild type. Wild-type plants showed an electrolyte leakage of 50% at −6.5 (±0.2, 95% CI) compared to *svk-1* (−7.7, ±0.2) and *uns-1* (−7.9, ±0.3). The increase in freezing tolerance is remarkable and is a similar to that of the *myb15* mutant, a transcription factor that directly suppresses *CBF* expression^24^. Together, our findings suggest that the genomic region encoding *SVK* represses cold-induced *CBF1* expression and freezing tolerance.

**Figure 2.**
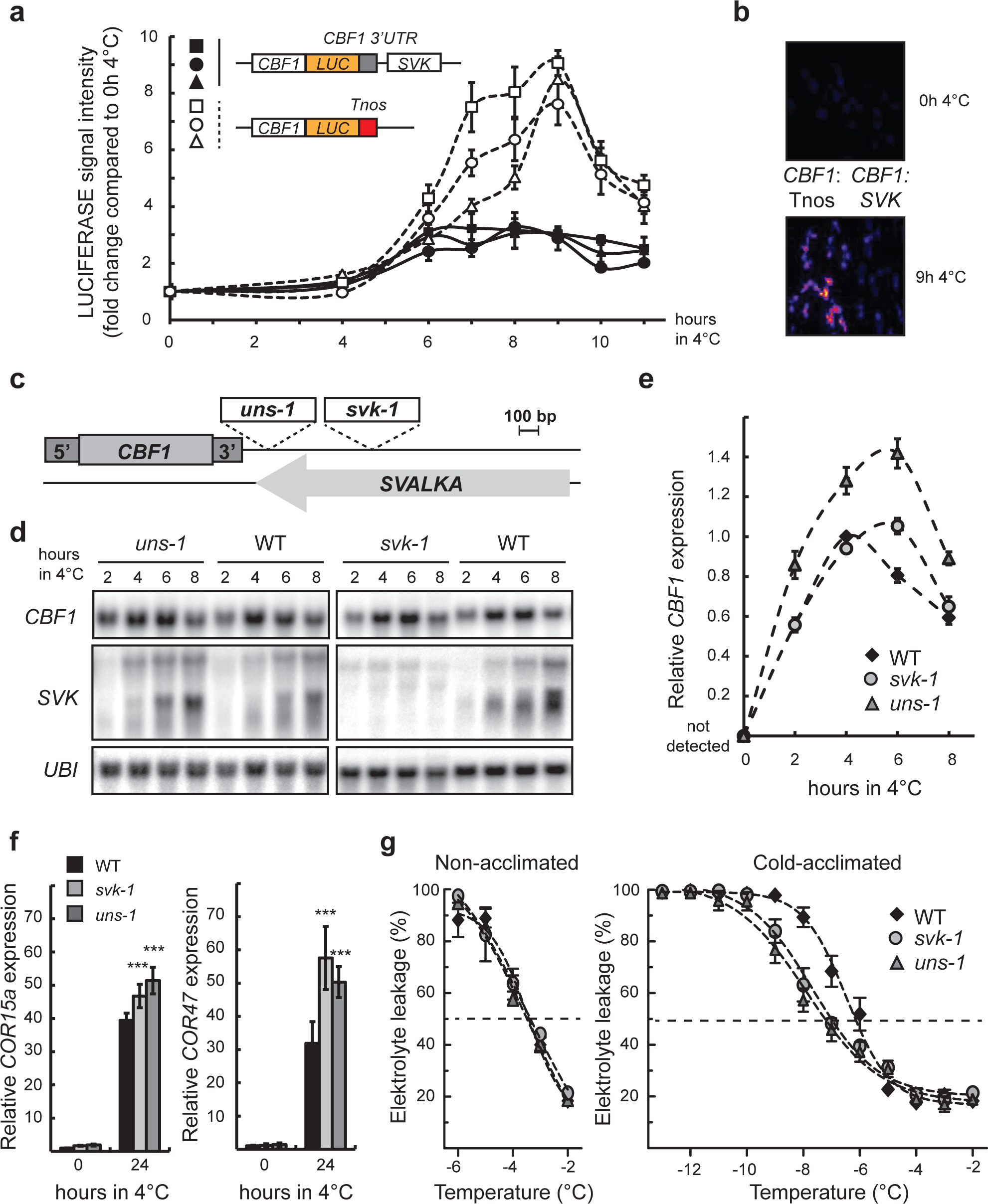
Characterization of the long non-coding RNA, *SVALKA*. a) CBF1-LUCIFERASE activity of constructs where *SVALKA* is present or not. White markers indicate three independent transformants of a construct where *SVALKA* is not present and black markers indicate three independent transformants of a constructs where *SVALKA* is present. Lines where *SVALKA* is present showed a decreased LUCIFERASE activity following cold exposure compared to lines *SVALKA* is not present. LUCIFERASE activity was measured as average pixel intensity of at least 5 seedlings for each time point. Markers represent mean with standard deviation. b) Representative images of lines in control conditions and after 9 hours of cold exposure for the two LUC constructs. c) Two T-DNA mutants disrupt the *CBF1-SVK* genomic region. The upper panel shows a graphical representation where the T-DNA insertions are located. d) A representative Northern blot of a cold exposure time series of Col-0 (WT), *uns-1* and *svk-1*. Probes for *CBF1* and *SVK* are the same as Figure 1. Blots were repeated with three biological replicates with similar results. *UBI* is used as loading control. e) Quantification of relative *CBF1* expression after different times of cold exposure in WT, *svk-1* and *uns-1*. Northern blot signal intensity from three biological replicates were normalized to their *UBI* signal and further normalized to the relative *CBF1* level in WT after 4h at 4°C. f) Relative level of *COR* transcripts in control and after 24h of cold exposure in WT, *svk-1* and *uns-1* determined by RT-qPCR. Bars represent mean (± standard deviation) from three biological replicates. The relative level of the *COR* transcripts were normalized to the level in WT in control conditions. Statistical significant differences were determined with Student’s t-test (***p<0.001). g) Electrolyte leakage in non-acclimated (left panel) and cold-acclimated (right panel) WT, *svk-1* and *uns-1* plants. Each marker represents the mean electrolyte leakage after freezing test compared to the total electrolyte content (± standard deviation) from at least three biological replicates.

### Read-through transcription of *SVK* results in *asCBF1*

To better understand the mechanism of *SVK* repression, we revisited the effects of the *uns-1* insertion. *uns-1* is inserted downstream of *SVK* yet results in equivalent cold-related defects compared to *svk-1*. These findings could be reconciled if *SVK* was promoting transcription of a cryptic antisense transcript into the *CBF1* gene body, since such a transcript might be disrupted in *uns-1*. To test this hypothesis, we carefully examined the presence of an antisense transcript mapping to the 3’-UTR of *CBF1* (Fig. 3a-b). *HEN2* is part of the nucleoplasmic 3’ to 5’ exosome responsible for degrading many types of non-coding RNA Polymerase II (RNAPII) transcripts^25^. We used a high resolution time series of cold exposed samples of wild type and *hen2-2* to enable the detection of cryptic antisense transcripts. A *CBF1* antisense transcript (*asCBF1*) corresponding to roughly 250 nt was detectable in the *hen2-2* mutant, yet not in the wild type (Fig. 3a, Supplementary Fig. 2a). *asCBF1* was also detected in additional mutants disrupting the nuclear exosome (Supplementary Fig. 2b). However, we could not detect *asCBF1* in the *sop1-5* (*suppressor of pas2*) mutant that have an increase of a subset of HEN2 dependent degradation targets^26^ or the *trl-1* mutant (Supplementary Fig. 2b). TRL1 (TRF4/5-like) is involved in 3’-processing of rRNA in the nucleolus^27^. Together, these results suggest *asCBF1* transcript is a nucleoplasmic exosome target with expression correlated to *SVK*. The size of *asCBF1* suggests it is either a *SVK* read-through transcript that is cleaved at the poly-(A) signal or a new transcription initiation event. To confirm that *asCBF1* depends on *SVK* transcription, we crossed the *svk-1* and *uns-1* mutants to *hen2-2* (Fig. 3c). In both double mutants *asCBF1* disappeared, confirming that *asCBF1* transcription depends on *SVK*.

**Figure 3.**
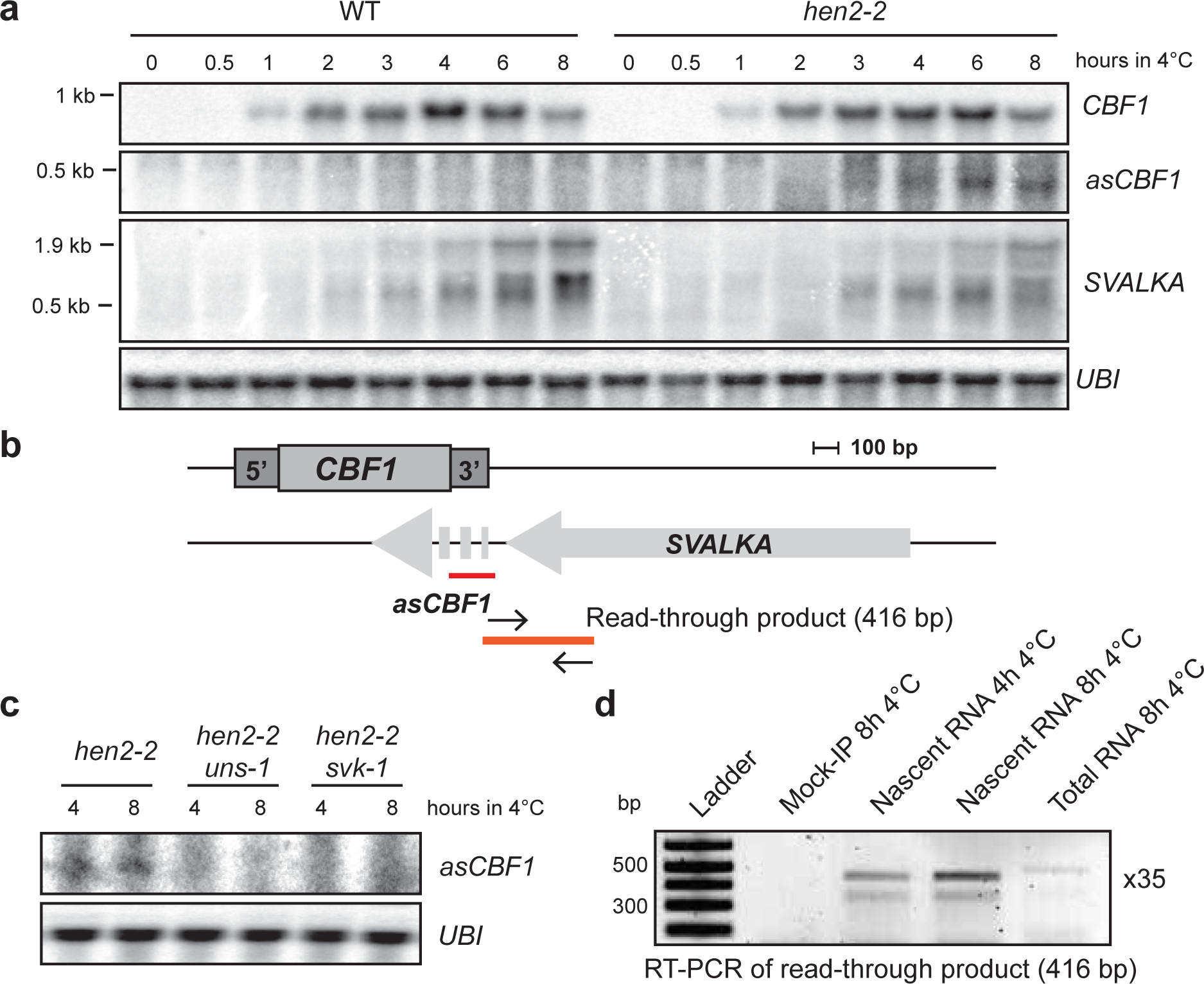
SVALKA transcription mediates transcription activity antisense to *CBF1*. a) Representative Northern blot of a cold exposure time series of Col-0 (WT) and *hen2-2*. Probes for *CBF1* and *SVK* are the same as Figure 2. The probe used for *asCBF1* is shown in B. Blots were repeated with three biological replicates with similar results. *UBI* is used as loading control. b) Graphical representation of the *CBF1-SVK* genomic region. The probe for *asCBF1* is shown in red and primers for the read-through transcript are shown by black arrows. Orange line indicates the read-through PCR product. c) Representative Northern blot of a cold exposure time series of *hen2-2* and the double mutants *hen2-2uns-1* and *hen2-2svk-1*. The probe used for *asCBF1* is shown in B. Blots were repeated with three biological replicates with similar results. UBI is used as loading control. d) RT-PCR of *SVALKA* read-through transcript. 100 ng of RNA from Pol II-IP or total RNA was converted to cDNA with a gene specific primer targeted to detect read-through *SVALKA* transcripts. A 416 bp read-through transcript can be detected in the nascent RNA samples with a more intense band after 8 hours of cold exposure. No PCR product could be seen in the Mock-IP or total RNA sample.

Transcription in higher eukaryotes occurs beyond the poly-(A) termination signal before nascent transcript cleavage is triggered^28^. Nascent transcript cleavage creates the 3’-end of the full length mRNA substrate for poly-adenylation, and a free 5’-end is attacked by 5’ to 3’ exonucleases eliciting transcription termination according to the torpedo model^29,30^. To detect any read-through transcription from *SVK*, we purified nascent RNA associating with RNAPII complexes (Supplementary Fig. 2c). We profiled this nascent RNA for the presence of *SVK* transcripts extending into *CBF1* as antisense transcripts (Fig. 3b, d). We detected a read-through PCR product after 4 hours of cold exposure and increased expression after 8 hours. Consistent with *asCBF1* being a product of read-through following *SVK* 3’end formation and cleavage, we could not detect the *SVK-asCBF1* read-through transcript in total RNA or in a Mock-IP control samples. Detection in nuclear exosome mutants and nascent transcript preparation gives *asCBF1* characteristics of a cryptic lncRNA that is quickly cleaved and degraded in wild-type. The XRN3 is thought to represent the 5’-to-3’ torpedo exonuclease mediating transcriptional termination in *Arabidopsis*^30,31^. However, we did not observe the *asCBF1* RNA or *CBF1* mis-regulation in the *xrn3-3* mutant, suggesting that this effect is XRN3 independent (Supplementary Fig. 2d). In summary, our findings revealed *asCBF1* as a *HEN2*-dependent “cryptic” antisense transcript that depends on *SVK* read-through transcription during cold to repress *CBF1* induction.

### *CBF1* expression is limited by RNAPII collision

Two molecular mechanisms could explain the observed *SVK*-mediated effects on *CBF1* expression: 1) transcription of *asCBF1* activates the siRNA pathway to trigger repression^32^, or 2) the act of *asCBF1* transcription in itself causes sense/antisense competition and RNAPII collision^33^. To test if siRNA were involved, we performed cold exposure of several different mutants in various siRNA pathways and probed for *CBF1* mRNA levels (Supplementary Fig. 3a). All the mutants failed to show increased *CBF1* levels (Supplementary Fig. 3b), ruling out the siRNA model. The RNAPII collision model predicts a discrepancy of transcription between the 5’-end and the 3’-end of *CBF1*. Colliding RNAPII complexes resulting from simultaneous sense/antisense transcription would result in pre-maturely terminated sense transcripts, and hence relatively more *5’-CBF1* transcripts than *3’-CBF1* transcripts. We tested these hypotheses using RNA samples taken after 4 and 8 hours of cold exposure (i.e. *asCBF1* independent versus *asCBF1* dependent time points). Northern blot analysis showed a peak of *CBF1* full length mRNA after 4 hours followed by a decrease after 8 hours (Fig. 4a). qPCR probes in the 5’-end of *CBF1* did not show a significant decrease between 4 and 8 hours at 4°C, while we detected a statistically significant difference with a probe specific to the *CBF1* 3’-end (Fig. 4b). These data suggest that the major decrease after 8 hours at 4°C in full length *CBF1* mRNA levels relate to differences in the 3’-end. Furthermore, we could detect a high amounts of pre-maturely terminated *CBF1* when sense transcripts are probed in the 5’-end compared to the 3’-end with Northern blot (Supplementary Fig. 4a). This effect was exaggerated in the *hen2-2* mutant, suggesting that many pre-terminated *CBF1* events are rapidly degraded by the nuclear exosome.

**Figure 4.**
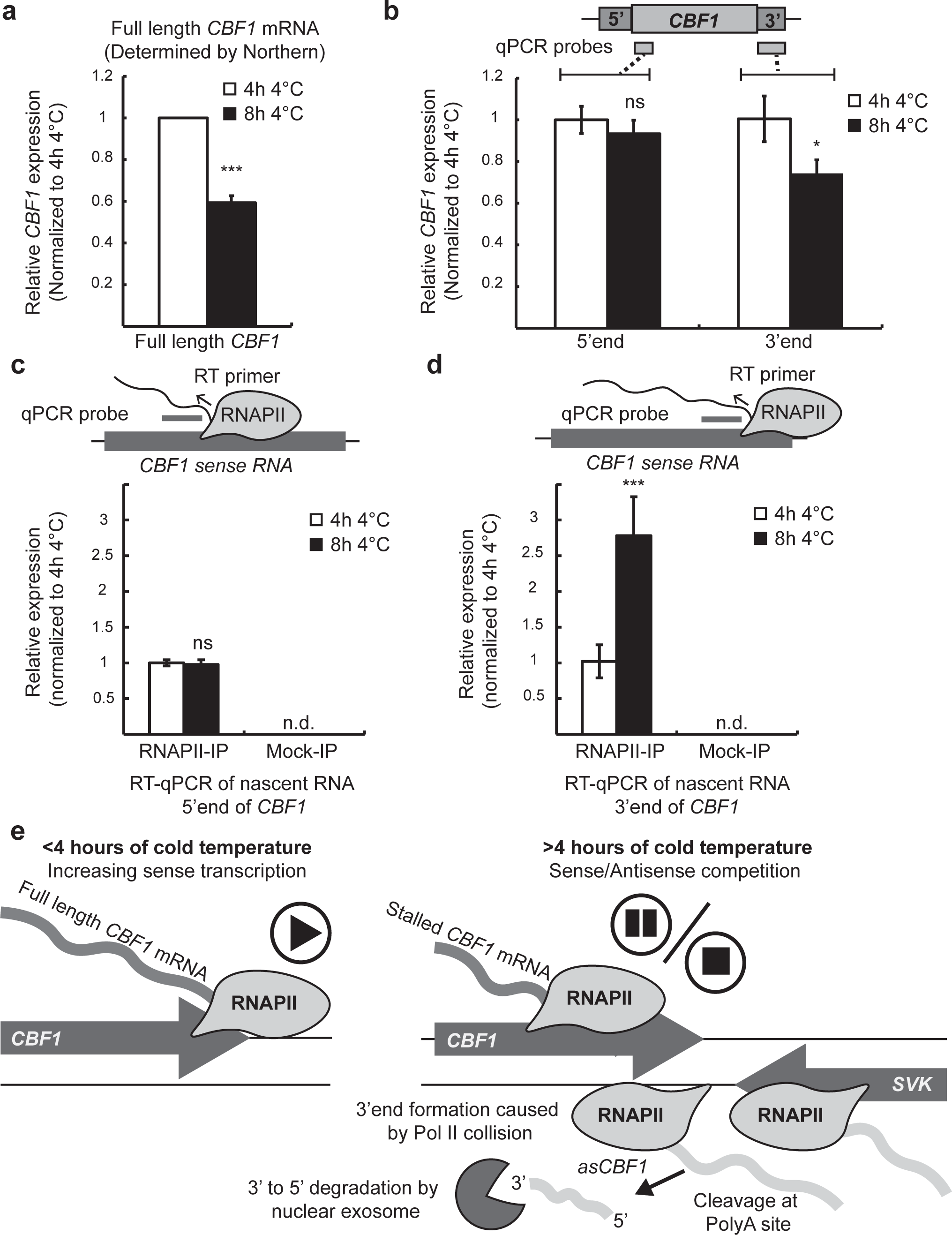
Antisense transcription to *CBF1* results in stalling mRNA transcription at the 3’end of *CBF1*. a) Quantification of full length *CBF1* transcript determined by Northern blots (Fig 2) using the probe shown in Figure 2. Graph represent mean from three biological replicates and relative to their loading control (UBI). b) RT-qPCR with probes in the 5’-end and 3’-end of *CBF1* using random primer generated cDNA as template. Bars represent mean (± standard deviation) from three biological replicates. The relative level of *CBF1* transcripts were normalized to the level at 4h 4°C for respective probe. Statistical significant differences were determined with Student’s t-test (*p<0.05). c) RT-qPCR with probe in the 5’-end of *CBF1* using nascent RNA converted into cDNA with gene specific primers as template. Pol II-IP was performed with a NRPB2-FLAG line and Anti-FLAG magnetic beads. Upper panel shows a graphical representation of the experimental setup. Bars in lower panel represent mean (± standard deviation) from three biological replicates. The relative level of *CBF1* transcripts were normalized to the level at 4h 4°C. Statistical significant differences were determined with Student’s t-test (ns, not significant). Mock-IP indicates RNA isolated from Col-0 (i.e. non-FLAG containing genotype). d) RT-qPCR with probe in the 3’-end of *CBF1* using nascent RNA converted into cDNA with gene specific primers as template. Pol II-IP was performed with a NRPB2-FLAG line and Anti-FLAG magnetic beads. Upper panel shows a graphical representation of the experimental setup. Bars in lower panel represent mean (± standard deviation) from three biological replicates. The relative level of *CBF1* transcripts were normalized to the level at 4h 4°C. Statistical significant differences were determined with Student’s t-test (***p<0.001). Mock-IP indicates RNA isolated from Col-0 (i.e. non-FLAG containing genotype). e) Representative Northern blot of a cold exposure time series of *hen2-2*, *cbf1-1* and the double mutant *hen2-2cbf1-1*. Blots were repeated with three biological replicates with similar results. *UBI* is used as loading control. f) Mechanistic model of how transcription of *SVK* represses sense *CBF1* transcription. Initially during cold exposure, SVK is not expressed and sense *CBF1* can transcribed without hindrance (upper panel). *CBF1* expression peaks after 4 hours of cold exposure. Simultaneously, expression of the lncRNA *SVK* is increased in the antisense direction of *CBF1* (lower panel). Read-through transcription of *SVK* results in transcription antisense to the 3’end of *CBF1* and an increase of Pol II occupancy on both strands. The intensification of transcribing polymerases in both directions creates Pol II collision and stalling of *CBF1* sense transcription. The outcome of the collision is that fewer Pol II complexes reach the end of the *CBF1* sense transcription unit and a decrease of full length *CBF1* mRNA.

An additional prediction of the RNAPII collision model involves the presence of stalled RNAPII transcription complexes in the 3’-end of *CBF1*^33^. Our protocol to isolate nascent RNA enabled us to detect strand-specific sites of RNAPII accumulation. The 5’-end probe showed no significant difference between 4 and 8 hours, suggesting that RNAPII occupancy early in *CBF1* elongation was indistinguishable (Fig. 4c). Intriguingly, the probe in the 3’-end showed that RNAPII occupancy was significantly increased over this region of *CBF1* after 8 h compared to 4 h (Fig. 4d). Collectively, these findings differ from the results assaying steady-state full length mRNA level (Fig. 4a). These results imply the presence of stalled RNAPII complexes over the 3’-end of *CBF1* that were correlated with increased *asCBF1* expression. In conclusion, our results showed that read-through of a lncRNA triggers head-to-head RNAPII collision over the *CBF1* gene body leading to regulated termination of *CBF1* transcription and a decrease of full length *CBF1* mRNA levels (Fig. 4e). This study illustrates how lncRNA read-through transcription limits peak expression of gene expression to promote an appropriate response to environmental change.

## Discussion

Environmental change alters RNAPII transcription in genomes so that organisms can adequately respond to new conditions^5,34^. These transcriptional responses include high RNAPII activity in non-coding regions, resulting in lncRNA transcription. On the one hand, lncRNA molecules may provide function, for example through riboswitch regulation^35,36^. On the other hand, the process of lncRNA transcription by RNAPII can affect the regulation of gene expression in the vicinity of target genes^37^. A curious result of widespread lncRNA transcription is that genes are often transcribed on both DNA strands, resulting in mRNA from the sense strand, and lncRNA from the antisense strand^38^.

Antisense transcription is detected in many genomes, and genome-wide analyses suggest that these phenomena can be either positively or negatively correlated with the corresponding sense transcript^39^. The positive correlations between sense and antisense expression represent a puzzling conundrum, as this relationship poses the question of how two converging RNAPII enzymes transcribing the same DNA can move past each other^33^. Sense and antisense transcription pairs on cell population level have been resolved as binary expression states for the individual transcripts at single-cell level^40^. Perhaps alternative expression states of sense or antisense lncRNA transcription in individual cells offers an explanation to why examples of gene repression by RNAPII collision are relatively rare. Overall, antisense transcription appears to be a common strategy to affect gene repression^39^.

Our study uncovers how the interplay between two lncRNA limits the production of maximal mRNA levels of the *Arabidopsis CBF1* gene by an RNAPII collision mechanism. Cold-induced transcription initiation in the *CBF* locus triggers expression of a cascade of antisense transcripts that fine-tunes the level of *CBF1* mRNA, thereby representing a negative feedback required for plants to appropriately acclimate to low temperatures.

The Identification of interplay between a stable lncRNA *SVALKA* and a cryptic lncRNA *asCBF1* that act together in gene repression represents an important aspect of our study. The cryptic lncRNA *asCBF1* is detectable after 6 hours of cold treatment in nuclear exosome mutant backgrounds. *asCBF1* is likely representative for additional lncRNA with regulatory role that are challenging to study owing to their cryptic and environmental specific expression characteristics. Our findings of multiple lncRNA acting together is line with recent findings suggesting a coordinated effect of multiple lncRNA to regulate meiosis in yeasts^41,42^. The combinations of multiple lncRNAs amplify the interfaces with regulatory potential and may increase precision of target regulation. Since *asCBF1* results from RNAPII read-through transcription of *SVK*, our findings highlight the process of transcriptional termination of lncRNA for regulation. Selective transcriptional termination of the *Arabidopsis* lncRNA *COOLAIR* determines *FLC* expression states by a chromatin-based mechanism^2^. Moreover, the class of budding yeast stable uncharacterized transcripts (SUTs) with environment-specific expression is characterized by inefficient termination resulting in read-through transcription into neighboring genes. Consistently, regulation of yeast gene expression by read-through transcription of SUTs is well-documented^43^. Extended SUT species link transcripts to create interdependency of loci to achieve coordinated expression changes^38^. The effects of the *SVK-asCBF1* circuitry support the idea that cascades of lncRNA transcription provide a mechanism to coordinate local regulation.

lncRNA cascades increase the spacing distance between initial lncRNA transcription event and the target gene, in contrast to a straightforward antisense lncRNA. Multiple cascading transcripts provide more room for independent regulatory inputs to achieve precision in regulation. The topology of the *Arabidopsis* genome is thought to be gene-centric with local gene loops rather than larger topologically associated domains (TADs)^44,45^. We would expect extensive co-regulation if sense and antisense lncRNA transcription were part of the same gene loop. Indeed, alternative gene loop formation is triggered by transcription of the lncRNA *APOLO* to regulate expression of the *PINOID* gene^46^. Read-through transcription of the lncRNA *SVALKA* may provide a mechanism to invade and break positive feedback mechanisms reinforcing *CBF1* transcription. As lncRNA transcription in yeast and mammalian cells correlate with the boundaries of chromatin domains, roles of lncRNA in communicating regulation across domain boundaries may be common despite differences in genomic topologies^47,48^.

The entire transcripts in the *CBF* cluster increase expression in response to short-term cold temperatures^11^. The discovery of the *SVK-asCBF1* lncRNA cascade illustrates how transcriptional activation of this region is associated with an inbuilt negative feedback mechanism that limits the timing of maximal *CBF1* expression by triggering RNAPII collision. While *CBF* genes promote freezing tolerance, excess *CBF* gene expression is associated with fitness penalties^16–19^. In this regard, the *SVK-asCBF1* cascade provides a mechanism to tightly control maximal *CBF1* expression and timing that could be exploited to engineer freezing tolerance with mitigated fitness costs. The conservation of *CBF* gene clusters and their role in promoting plant freezing tolerance suggests that lncRNA cascades equivalent to *SVK-asCBF1* exist in other species. Our research suggests that the cryptic lncRNA component in such cascades could be identified by transcript profiling in nuclear exosome mutants. The lncRNA-mediated effects of long-term cold exposure on plant gene expression have been well characterized for the *FLC* gene^2,40^. Our research demonstrates that lncRNAs also mediate the response to short-term and transient cold exposure by timing maximal gene expression. Future research will be needed to investigate if biophysical properties of lncRNA transcription may provide a particularly effective sensor for temperature. Perhaps more likely, lncRNA functions in temperature responses coincide with the intensity of research efforts addressing this question in plants to prepare for changing climates. In sum, our findings provide a compelling example of functional lncRNA transcription in cells to limit gene induction by triggering RNAPII collisions.

## Methods

### Plant materials and growth conditions

*A. thaliana* seedlings were grown on ½ MS +1% Sucrose plates in long day photoperiod (16 h light/8 h dark) at 22°C/18°C unless otherwise stated. For all experiments, Col-0 was used as wild-type background. Genotypes used in this study can be found in supplementary table 1. For cold treatment, seedlings were grown in control conditions for 10 days (100 μE⋅m^−2^⋅s^−1^) and subsequently transferred to 4°C for indicated times and sampled. Light intensity during cold treatment was approximately 25 μE⋅m^−2^⋅s^−1^. For freezing tests, plants were grown in short day conditions (8 h light/16 h dark) at 22°C/18°C for 6 weeks. Cold acclimation was performed in short day conditions at 4°C for 4 days prior to freezing test.

### Cloning and Luciferase assay

Primers for cloning can be found in Supplementary table 2. For the CBF1:LUC:Tnos construct, a fragment encompassing the *CBF1* promoter and gene body (−1903 bp to +639 bp relative to the translation start) was amplified from genomic DNA with Phusion polymerase (Thermo Fisher Scientific, USA) and ligated to the pENTR/D-TOPO vector (Invitrogen, USA). After sequencing to eliminate any cloning errors the pENTR/CBF1 vector was incubated with pGWB535 vector in a LR reaction. The final vector, pGWB535/CBF1:LUC:Tnos was transformed into Col-0 plants using the floral dip method (Clough & Bent, 1998). For the CBF1:LUC:SVK construct, a fragment encompassing the *SVK* promoter and transcription unit (+643 bp to +3410 bp relative to the *CBF1* translation start) was amplified from genomic DNA and fused to a CBF1:LUC fragment amplified from pGWB535/CBF1:LUC:Tnos. The fused fragment was ligated to pENTR/D-TOPO and sequenced. A LR reaction between pENTR/CBF1:LUC:SVK and pGWB501 produced the final vector, PGWB501/CBF1:LUC:SVK, which was transformed into Col-0. Transformed seedlings were selected on Hygromycin plates and their progeny (T2 generation) was used for the Luciferase assay. For detection of LUC, 10 day old seedlings were treated with cold temperature (4°C). At indicated times 5 µM D-Luciferin (ZellBio, Germany) was sprayed onto seedlings followed by dark incubation for 30-60 min. LUC was detected using a CCD camera and pixel intensity was determined in ImageJ. The average pixel intensity from at least 5 seedlings was used to get a mean value and used for statistical analysis.

### RNA extraction, Northern analysis, 5’- and 3’-RACE and RT-qPCR

Total RNA was extracted using RNeasy Plant Mini Kit (Qiagen, Germany) according to manufacturer’s instructions. Northern analysis was performed as described with minor modifications^43^. In short, 5-20 µg of total RNA was separated on a 1.2% agarose gel with formaldehyde and 1xMOPS. Gels were blotted overnight onto a nylon membrane and crosslinked with UV radiation. Probes were made in a PCR reaction by incorporating radioactive dTTP (^32^P, PerkinElmer, USA) from a DNA probe template. Membranes were exposed to a phosphorimager screen (GE Healthcare, UK) for 1-10 days depending on the expression level of the transcript of interest. Screen was subsequently scanned with a Typhoon scanner (GE Healthcare, UK). RACE experiments were done with a SMARTer RACE 5’/3’ Kit (Takara, Japan) according to manufacturer’s instructions. For RT-qPCR, total RNA was DNase treated with TURBO DNase (Thermo Fisher Scientific, USA) and 1 µg DNase treated RNA was subsequently turned into cDNA as per manufacturer’s instructions with Superscript III (Invitrogen, USA) using random primers, oligo dT or gene specific primers depending on the experiment. Diluted cDNA (1:10) was used in a PCR reaction with GoTaq qPCR Master mix (Promega, USA) and run on a CFX384 Touch instrument (Bio-Rad, USA). Data was processed in CFX manager and exported to Excel (Microsoft, USA) for further analysis. Relative expression was calculated and normalized to at least two internal reference genes. All primers used in this study can be found in Supplementary table 2.

### Freezing test

Freezing test was performed as described^49^. Briefly, leaf discs of non-acclimated or cold-acclimated plants were put in glass tubes with 200 μl of deionized water. The tubes were then transferred to a programmable bath at −2°C (FP51, Julabo, Germany). After 1 hour, ice formation was induced and the temperature was slowly decreased (−2°C/h). Samples were taken out of the bath at designated temperatures and cooled on ice for an hour followed by 4°C. When all samples were collected, 1.3 ml of deionized water was added and the tubes were shaken overnight at 4°C. Electrolyte leakage was measured using a conductivity cell (CDM210, Radiometer, Denmark). To get total ion content, tubes were immersed in liquid nitrogen, thawed, shaken again overnight and measured for conductivity. Electrolyte leakage was determined by comparing the measured conductivity before and after the liquid nitrogen treatment. Data was fitted to a sigmoidal dose-response with GraphPad Prism.

### TSS-seq and bioinformatic analysis

The genome-wide distribution of TSSs in wild-type was recently mapped in *Arabidopsis* using 5’-CAP-sequencing^50^. Here, we extend these analyses to 2-week old cold-treated seedlings (3 hours at 4°C). 5 micrograms of DNase-treated total RNA were treated with CIP (NEB) to remove non-capped RNA species. 5’-caps were removed using Cap-Clip (CellScript) to permit ligation of single-stranded rP5_RND adapter to 5’-ends with T4 RNA ligase 1 (NEB). Poly(A)-enriched RNAs were captured with oligo(dT) Dynabeads (Thermo Fisher Scientific) according to manufacturer’s instructions and fragmented in fragmentation buffer (50 mM Tris acetate pH 8.1, 100 mM KOAc, 30 mM MgOA) for 5 mins at 80°C. First-strand cDNA was generated using SuperScript III (Invitrogen) and random primers following manufacturer’s instructions. Second-strand cDNA was generated with the BioNotI-P5-PET oligo and using Phusion High-Fidelity Polymerase (NEB) as per manufacturer’s instructions. Biotinylated PCR products were captured by streptavidin-coupled Dynabeads (Thermo Fisher Scientific), end repaired with End Repair Enzyme mix (NEB), A-tailed with Klenow fragment exo-(NEB), and ligated to barcoded Illumina compatible adapter using T4 DNA ligase (NEB). Libraries were amplified by PCR, size selected using AMPure XP beads (Beckman Coulter), pooled following quantification by bioanalyzer, and sequenced in single end mode on the following flowcell: NextSeq® 500/550 High Output Kit v2 (75 cycles) (Illumina). For the bioinformatics, all supplementary code for the data analysis pipelines described below is available at https://github.com/Maxim-Ivanov/Kindgren_et_al_2018. The NGS data manipulations were detailed in the 01-Processing_5Cap-Seq_data.sh pipeline. In brief, the custom adapter sequences (ATCTCGTATGCCG) were trimmed from 3’ ends of the 5Cap-Seq (TSS-Seq) reads using Trim Galore v0.4.3. Then 8 nt random barcodes (unique molecular identifiers, or UMIs) were trimmed from 5’ ends of reads and appended to read names using a custom script (UMI_to_Fastq_header.py). The resultant Fastq files were aligned to TAIR10 genome using STAR v2.5.2b in the end-to-end mode^51^. SAM files were sorted and converted to BAM using Samtools v1.7^52^. Reads overlapping the rRNA, tRNA, snRNA and snoRNA genes (obtained from the Araport11 annotation) were filtered out using Bedtools v2.25.0^53^. In addition, multimapper reads with MAPQ score below 10 were removed by Samtools. Morever, we filtered out PCR duplicates by analyzing groups of reads sharing the same 5’ genomic position and removing reads with redundant UMIs (this was done using a custom script Deduplicate_BAM_files_on_UMI.py). Finally, stranded Bedgraph files were generated using Bedtools. For visualization in genomic browsers, the forward and reverse Bedgraph files corresponding to the same sample were combined together and normalized to 1 million tags. For the downstream analysis in R environment, the stranded Bedgraph files were filtered for coverage ≥ 2x and ensured to contain only genomic intervals with 1 bp width (Expand_bedGraph_to_single_base_resolution.py). The detection of 5’ tag clusters (TCs) was described in detail in the 02-Calling_5Cap-Seq_TSS.R pipeline. It makes an extensive use of the CAGEfightR package (https://github.com/MalteThodberg/CAGEfightR). Differentially expressed TCs were called using the DESeq2 package.

### Polymerase II immunoprecipitation and nascent RNA purification

5 grams of seedlings from a line where a NRPB2-FLAG construct covers a lethal *nrpb2-1* mutation (described in^54^) were grinded to a fine powder. For Mock-IP 5 grams of Col-0 seedlings were used. 15 ml of ice-cold extraction buffer (20 mM Tris-HCl pH 7.5, 300 mM NaCl, 5 mM MgCl_2_ (+ 5 µl/ml 20% Tween, 1 µl/ml RNAseOUT, 5 µl/ml 1 M DTT, prot. inhibitor tablet (Roche, 1 tablet for 50 ml buffer)) was added to the powder and 660 U of DNase I was subsequently added to the mix. After 20 minutes on ice, the mixture was put into a centrifuge for 10 min (4°C, 10000 g) and the supernatant was recovered. The supernatant was filtered through a 0.45 µm filter and added to Anti-FLAG M2 magnetic beads (Sigma-Aldrich, USA). Supernatant and beads were then incubated on a slowly rotating wheel at 4°C for 3 hours. Beads were washed 8 times with extraction buffer and immunoprecipitated polymerase complexes were eluted with 2 mg/ml 3xFLAG peptide (ApexBio, USA) for 2 times 30 minutes. Purified polymerase complexes was disrupted and attached RNA was extracted by the miRNeasy Mini Kit (Qiagen, Germany) according to manufacturer’s instructions. 100 ng isolated RNA was used to create cDNA with Superscript III (Invitrogen, USA) and gene specific primers.

### Western blotting

Protein samples were separated on a 4-15% TGX gel (Bio-Rad, USA) and blotted onto a PVDF membrane. Membranes were blocked in 5% milk powder dissolved in PBS. Primary antibody was incubated with membrane overnight at 4°C. Following morning, membranes were washed in PBS+T 2 times 10 min and incubated with secondary antibody (Agilent, USA) for 60 min at room temperature. After 3 times 5 min washes in PBS+T proteins were detected with Super-Signal West Pico Chemiluminescent (Thermo Fisher Scientific, USA) and developed with a ChemiDoc MP instrument (Bio-Rad, USA). Following antibodies were used in this study: Monoclonal Anti-FLAG M2 antibody (Sigma-Aldrich, USA) and Anti-RBP1 antibody (ab140509, abcam, UK).

## Acknowledgements

We would like to thank Prof. Åsa Strand and Sofie Grönlund for assistance with the freezing test. We thank Prof. Peter Brodersen for sharing seed and assistance with the Luciferase assay. We thank Prof. Vicent Pelechano for help with the 5’CAP-seq. We thank Jasmin Dilgen for technical assistance, Jan Høstrup for plant care and members of the S.M. laboratory for critical reading of the manuscript. Research in the laboratory of S.M. is supported by a Hallas-Møller Investigator award by the Novo Nordisk Foundation NNF15OC0014202, a Copenhagen Plant Science Centre Young Investigator Starting grant and the ERC StG2017-757411. R.A. is supported by an EMBO LTF (ALTF 463-2016). P.K. is supported by MSCA-IF 703085.

## Author contributions

P.K., R.A. and M.I. performed the experiments. P.K., R.A. and M.I. analyzed the data. S.M. supervised the project. P.K. and S.M. wrote the manuscript.

---

**Supplementary Figure 1.**
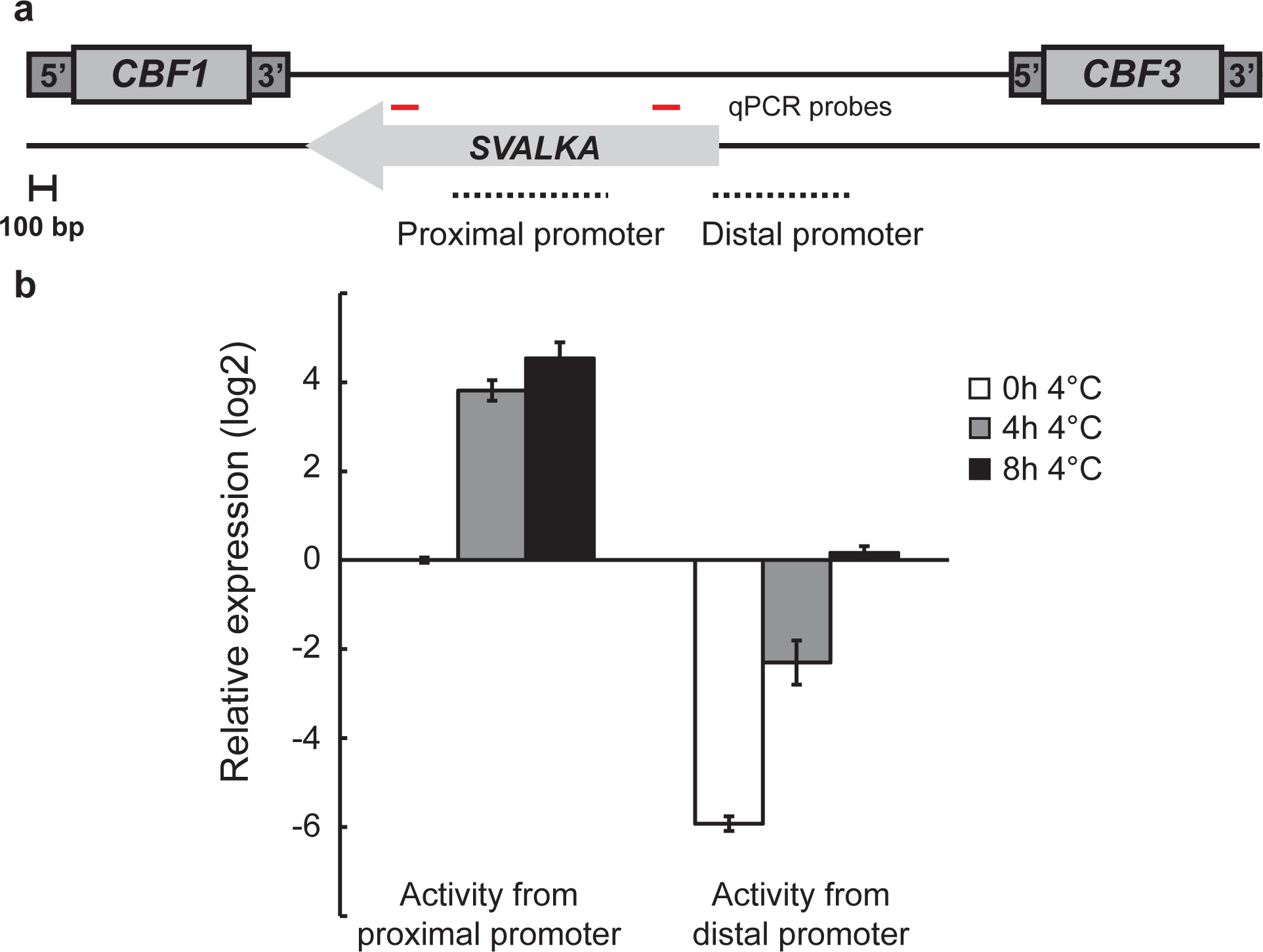
Promoter usage of *SVK*. a) Graphical representation of the *CBF1-SVK* genomic region. The red lines indicate the position of the qPCR probes in B. b) RT-qPCR with probes detecting transcripts from the proximal and distal promoter of *SVK*. Bars represent mean (± standard deviation) from three biological replicates. The relative level of *SVK* transcripts were normalized to the level at 0h 4°C for the proximal probe.

**Supplementary Figure 2.**
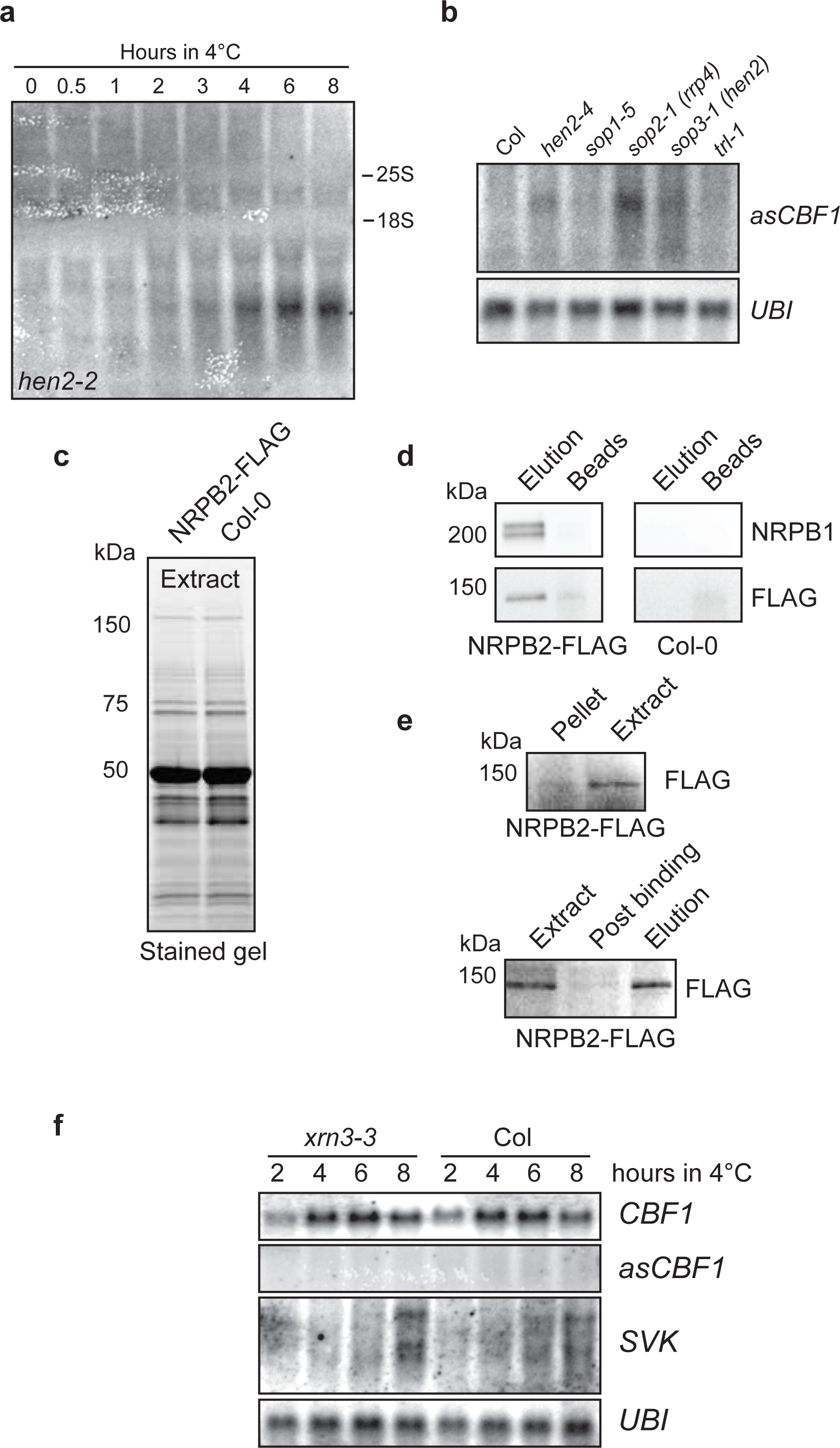
Purification of nascent RNA, characterization and mutant analysis of asCBF1. a) Representative Northern blot of a cold exposure time series of *hen2-2*. Blots were repeated with three biological replicates with similar results. Blot depicts the full membrane of the cut image shown in Figure 4A. b) Representative Northern blot of cold exposure (4h 4°C) of nuclear exosome mutants. Blots were repeated with three biological replicates with similar results. *UBI* is used as loading control. c) Input material for purification of Pol II complexes visualized from a stain-free TGX-gel. d) Western blots of elution from magnetic beads with Anti-FLAG and Anti-NRBP1 antibodies. From the NRPB2-FLAG line, both FLAG and NRBP1 were eluted indicating that Pol II complexes were still assembled in the elution. Most of the Pol II complexes bound to beads were eluted since no signal was detected from the remaining bead fraction. No signal could be detected in the non-FLAG genotype (Col-0). e) Western blots to control the purification of Pol II complexes monitored with Anti-FLAG. Most NRBP2-FLAG signal was found in the extract (upper panel) and eluted from the beads (lower panel). f) Representative Northern blot of a cold exposure time series of WT and *xrn3-3*. Blots were repeated with three biological replicates with similar results. *UBI* is used as loading control.

**Supplementary Figure 3.**
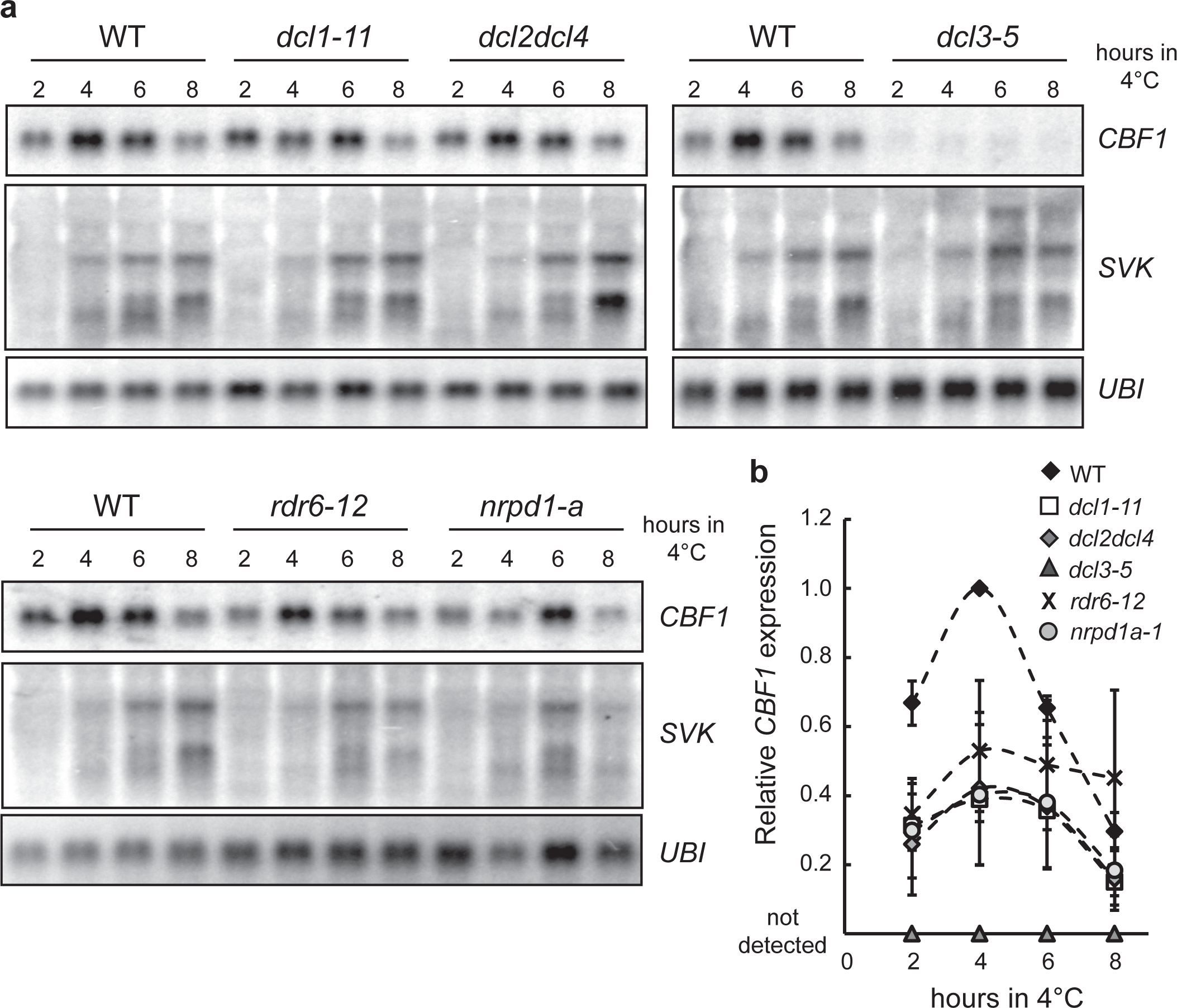
siRNA is not involved in *SVK*-mediated repression of *CBF1*. a) Representative Northern blots of a cold exposure time series of mutants in different siRNA pathways. Blots were repeated with three biological replicates with similar results. *UBI* is used as loading control. b) Quantification of relative *CBF1* expression after different times of cold exposure in WT and siRNA mutants. Northern blot signal intensity from three biological replicates were normalized to their *UBI* signal and further normalized to the relative *CBF1* level in WT after 4h at 4°C.

**Supplementary Figure 4.**
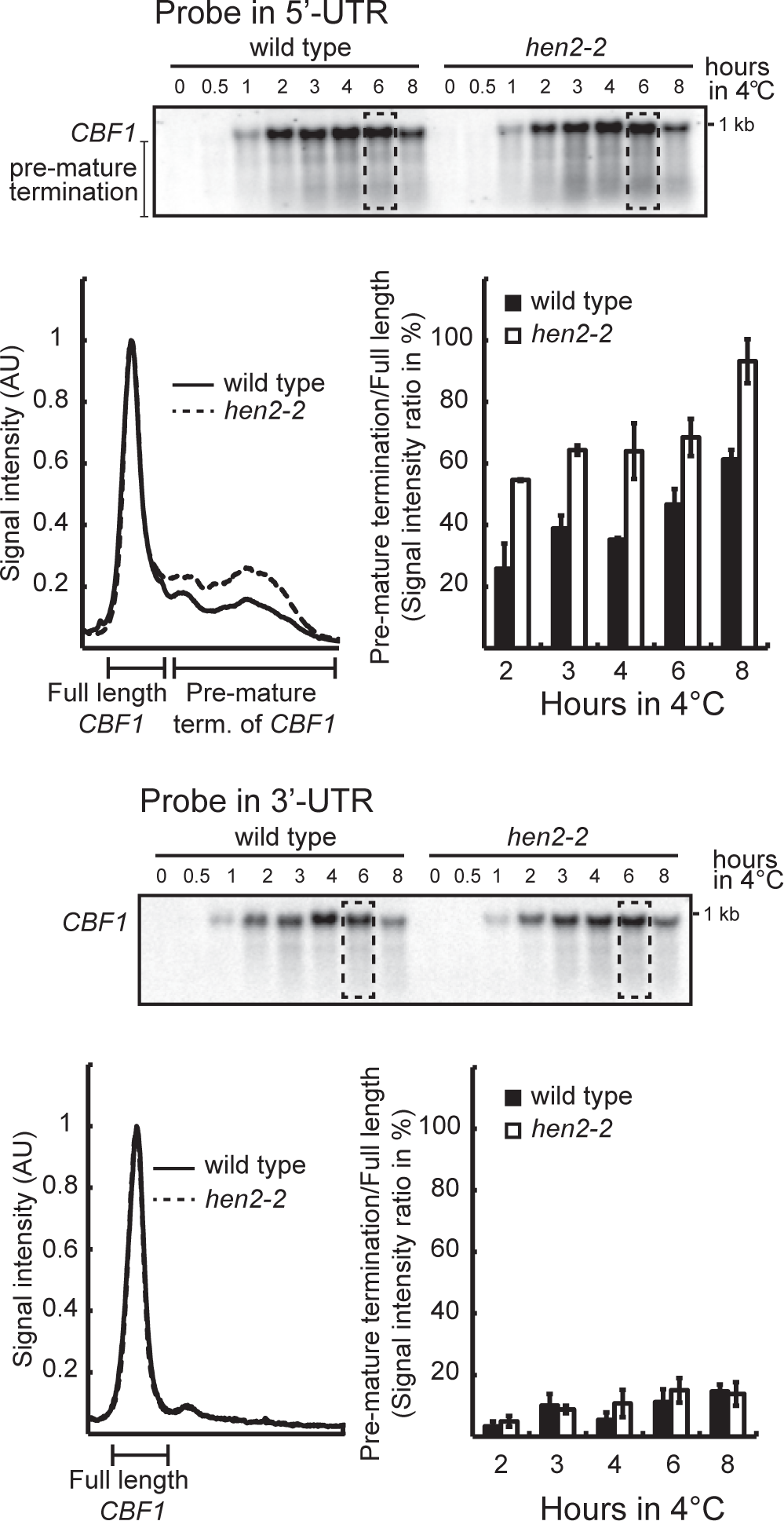
Characterization of the mechanism of *SVK*-mediated repression of *CBF1*. a) Detection of pre-maturely terminated *CBF1* determined by Northern blot in Col-0 and *hen2-2*. A probe in the 5’-UTR of *CBF1* detected an increased fraction of pre-terminated CBF1 (upper panel). A visualization of the signal intensity across the boxed square indicated a higher fraction of pre-terminated *CBF1* in the *hen2-2* mutant. Quantification of all lanes with three biological replicates showed an increase of terminated *CBF1* with increased duration of cold exposure. The combined signal from pre-terminated *CBF1* was represented as a percentage to the signal from full length *CBF1* for each time point. At all time-points the signal from *hen2-2* was higher than Col-0. A probe in the 3’-UTR showed small differences between Col-0 and *hen2-2* (lower panel).

**Supplementary table 1. Significantly up- and down-regulated TSS peaks in response to 3 of hours of 4°C.**

**Supplementary table 2. Genotypes used in this study.**

**Supplementary table 3. Primers used in this study.**

